# Single-cell transcriptomics of resected human traumatic brain injury tissues reveals acute activation of endogenous retroviruses in oligodendroglia

**DOI:** 10.1101/2022.09.07.506982

**Authors:** Raquel Garza, Yogita Sharma, Diahann Atacho, Arun Thiruvalluvan, Sami Abu Hamdeh, Marie Jönsson, Vivien Horvath, Anita Adami, Martin Ingelsson, Patric Jern, Molly Gale Hammell, Elisabet Englund, Agnete Kirkeby, Johan Jakobsson, Niklas Marklund

**Author notes:** Correspondence to: Niklas Marklund, Department of Clinical Sciences Lund, Neurosurgery, Lund University, Skåne University Hospital, Skåne University Hospital EA-blocket plan 4, Klinikgatan 17A, 222 42 Lund, Sweden. Johan Jakobsson, Lund Stem Cell Center, Department of Experimental Medical Sciences, Laboratory of Molecular Neurogenetics, Lund University, Sölvegatan 17, BMC A11, 221 84 Lund, Sweden. Co-shared senior authorship (Johan Jakobsson, Niklas Marklund).

## Abstract

Traumatic brain injury (TBI) is a leading cause of chronic brain impairment and results in a robust, but poorly understood, neuroinflammatory response that contributes to the long-term pathology. We used snRNA-seq to study transcriptomic changes in different cell populations in human brain tissue obtained acutely after severe, life-threatening TBI. This revealed a unique transcriptional response in oligodendrocyte precursors and mature oligodendrocytes, including the activation of a robust innate immune response, indicating an important role for oligodendroglia in the initiation of neuroinflammation. The activation of an innate immune response correlated with transcriptional upregulation of endogenous retroviruses in oligodendroglia. This observation was causally linked *in vitro* using human glial progenitors, implicating these ancient viral sequences in human neuroinflammation. In summary, this work provides a unique insight into the initiating events of the neuroinflammatory response in TBI, which has new therapeutic implications.

## Introduction

Traumatic brain injury (TBI) caused by, for example, motor vehicle accidents, violence or falls is a leading cause of mortality and persisting morbidity. The impact to the head results in immediate death to neuronal and glial cells, as well as injury to axons and the microvasculature. This initial primary brain injury is then substantially exacerbated by a poorly understood and progressive cascade of secondary events that may ultimately result in severe long-term consequences. This chronic phase of the injury is often characterized by persistent white matter atrophy, linked to neuroinflammation^1, 2^, and an increased risk of neurodegenerative disorders, such as Alzheimer’s or Parkinson’s disease ^3, 4^.

Clinical and experimental evidence suggests that a robust neuroinflammatory response plays a key role in the subsequent development of posttraumatic disability and the increased risk of chronic neurodegenerative disorders ^2, 5–7^. The neuroinflammatory cascade after TBI is complex and involves the activation of resident microglia as well the recruitment of peripheral immune cells. In addition, other resident brain cells such as astrocytes, oligodendrocyte precursor cells (OPCs) and mature oligodendrocytes may be directly involved in modulating or driving the neuroinflammatory response, although their role remains poorly understood ^8–10^.

Myelinating oligodendrocytes are essential for saltatory nerve conduction in the central nervous system and are involved in the metabolic support of neurons and the modulation of neuronal excitability ^11^. There is also emerging evidence that brain oligodendrocytes are morphologically and transcriptionally heterogeneous, and that their functional role may be altered in disease ^12, 13^. The neuroinflammatory response occurring after TBI seems to be associated with oligodendroglia vulnerability ^9^, and persistent neuroinflammation is often found in atrophied white matter tracts of long-term TBI survivors ^2^. These observations argue for an important contribution of the neuroinflammatory response to white matter injury. Notably, OPCs and oligodendrocytes have recently been shown to activate the transcription of immune-response genes in certain disease contexts ^14, 15^, suggesting that these cells may play a direct role in the neuroinflammatory process by adopting an immune-like cell state. Despite these observations, the role of oligodendroglia in the neuroinflammatory response occurring after TBI remains poorly understood.

While the presence of neuroinflammation after TBI, and in other neurodegenerative conditions, has been firmly established, the molecular mechanisms contributing to this sterile inflammatory response remains largely unknown. We and others have recently found that transcriptional activation of endogenous retroviruses (ERVs) or other transposable elements (TEs) are involved in the neuroinflammatory response in mouse models ^16, 17^. ERVs are remnants of old retrovirus infections that have entered our germline and make up about 8% of our genome. Recent evidence suggests ERVs can be transcriptionally activated in certain neurological disease states ^16^. This aberrant expression of ERVs results in the formation of double-stranded RNAs, reverse-transcribed DNA molecules, and ERV-derived peptides that induce a “viral mimicry”, where cells of the central nervous system respond by activating the innate immune system in the form of an interferon response, as if they were infected ^18–20^. This event results in a downstream trigger or boost of inflammation that may participate in the respective disease processes. If and how ERVs are activated in human brain disorders and after insults, such as in TBI, remains poorly documented.

In this study, we performed single nuclei RNA sequencing (snRNA-seq) on fresh-frozen brain tissue, which had been surgically removed due to life-threatening TBI. We found that TBI resulted in a unique transcriptional response in several cell types in the injured human brain tissue, including the activation of cell-cycle-related genes in microglia as well as alterations in synaptic gene expression in excitatory neurons. Notably, we detected clear evidence of an innate immune response in both OPCs and oligodendrocytes, including evidence for an interferon response. This innate immune response was accompanied with the transcriptional activation of the major histocompatibility complex (MHC) class I and II. These results demonstrate that oligodendroglia undergo a transformation to an immune-like cell state after TBI and suggest a key role for these cells in the initiation of neuroinflammation following such an insult. Moreover, the activation of the innate immune response was linked to transcriptional activation of ERVs in OPCs and oligodendrocytes, a phenomenon that could be modeled and replicated in human glial progenitors *in vitro*, implicating a potential role for these ancient viral sequences in human neuroinflammation. These results not only provide novel insights into initiating events of the neuroinflammatory response following severe TBI, but also opens up new therapeutic avenues.

## Results

### Samples and experimental design

To investigate cell-type-specific transcriptional responses after severe, acute TBI, we performed single-nuclei RNA sequencing (snRNA-seq) from fresh-frozen human brain tissue (Fig 1a). We recruited 12 severe TBI patients, defined as post-resuscitation Glasgow Coma Scale score ≤ 8. Detailed demographic and clinical characteristics are shown in Table 1. The age (mean ± standard deviation) of the patients (10 males, 2 females) was 49.5 ± 18.2 years. In these patients, decompressive surgery was a life-saving measure to remove injured and swollen space-occupying brain tissue causing marked mass effect or increased intracranial pressure (ICP) refractory to conservative, medical neurointensive care treatment. The injured and contused brain regions (typically the injured part of a temporal or frontal lobe) were surgically removed between 4 hours and 8 days after injury ^21^. As control tissue, we used five fresh-frozen *post-mortem* samples from the frontal and temporal lobe obtained from three non-neurological deaths of patients aged 69, 75 and 87 years (Table 1).

**Figure 1.**
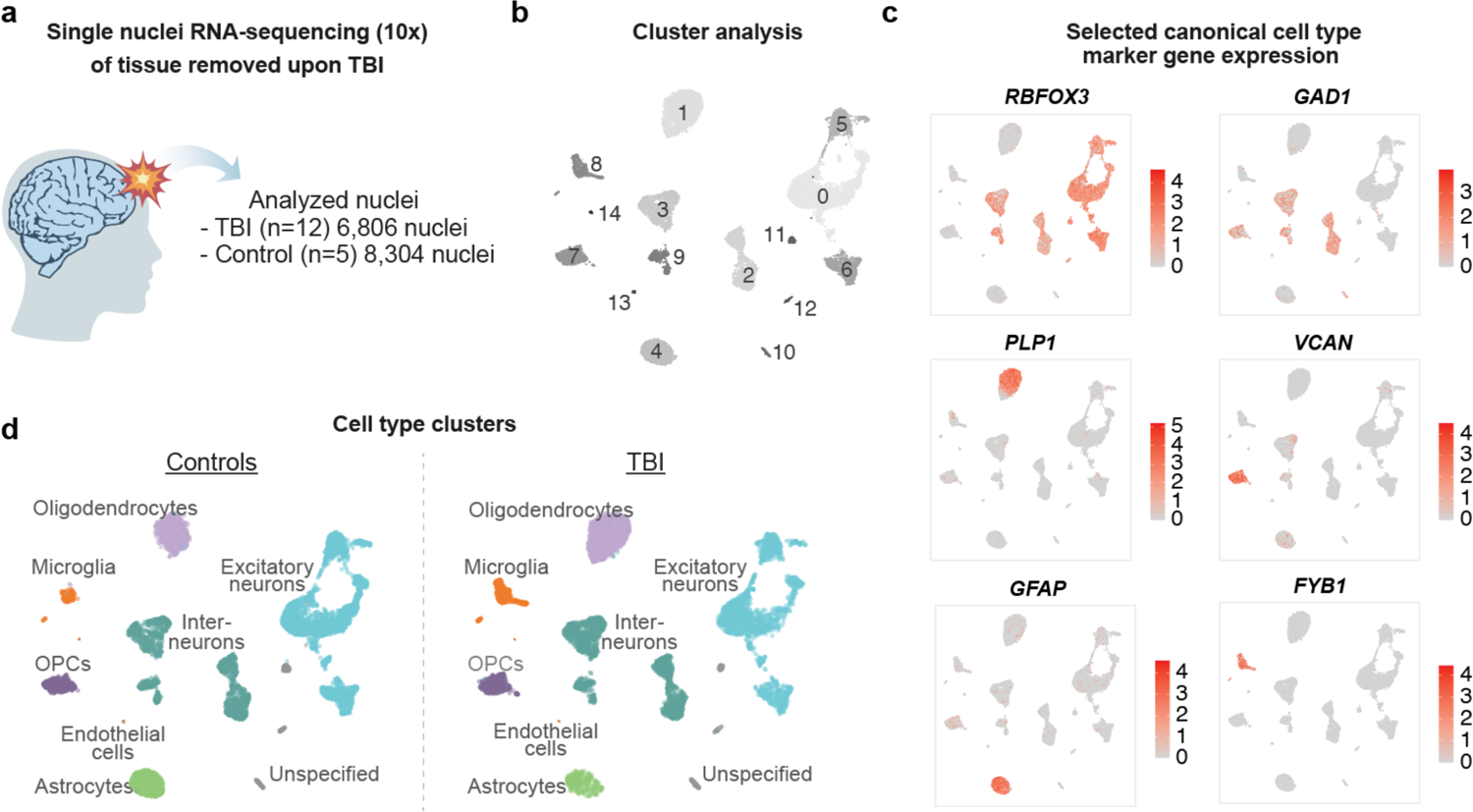
a) Schematic of experimental approach. Brain tissue was surgically removed after severe TBI followed by single-nuclei RNA-seq. Numbers indicate the number of nuclei recovered after quality control per condition. b) UMAP labelled with the nuclei clusters identified (clusters 0-14). c) Projection of gene expression of canonical gene markers to identify cell types. d) UMAP labelled by characterized cell types, split by condition.

**Table 1.**
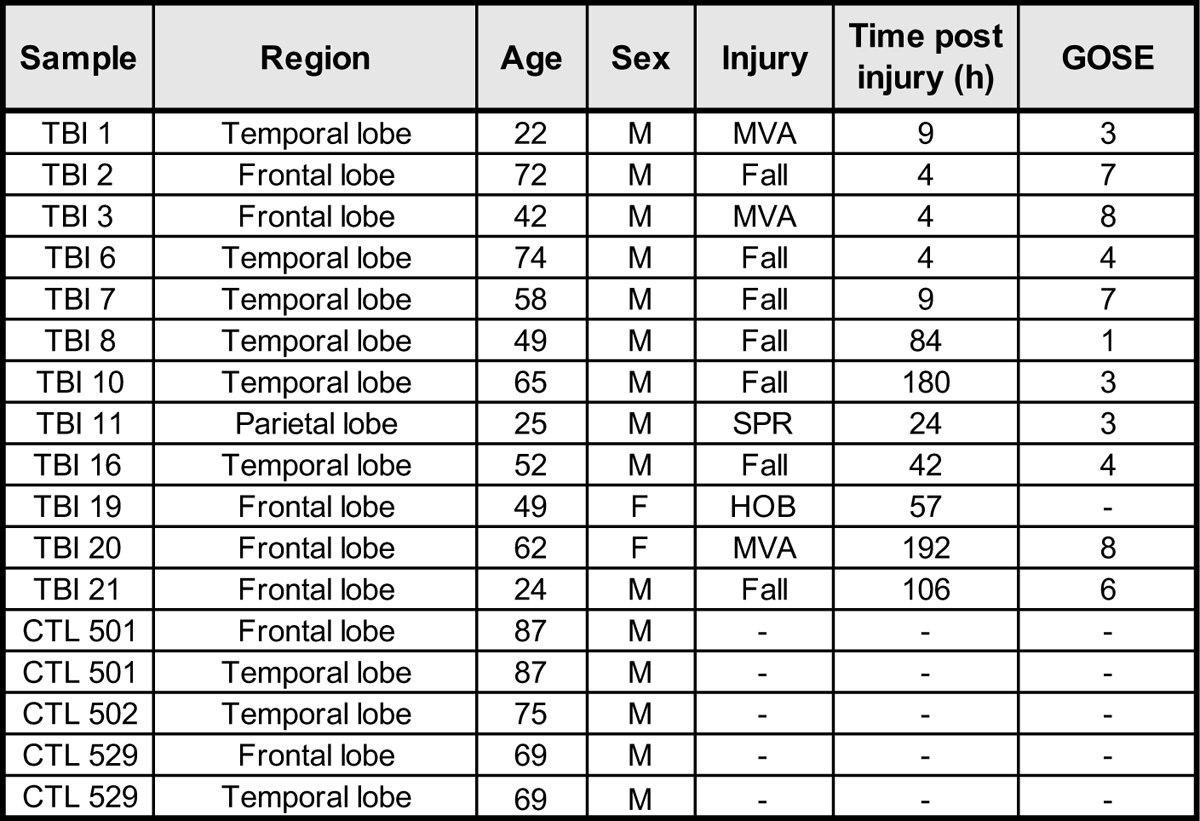
Demographics of TBI cases and controls, including the brain region of sample collection. MVA = motor vehicle accident, SPR = sports related, HOB = hit by object. GOSE = Glasgow Outcome Scale extended

We isolated nuclei from frozen human brain tissue and performed snRNA-seq using the 10X Genomics pipeline. The tissue obtained from TBI patients contained a high degree of blood and damaged cells, which greatly influenced the quality of sequencing data. Given the quality differences between the control and TBI samples, we used a strict threshold to select cells from both conditions to avoid bias due to differences in sequencing depth: only cells with at least 1,000 detected genes were used for further analysis (Suppl Fig 1a). After quality control (QC) ^22^, we kept high-quality sequencing data from the 12 TBI samples, with a total of 6,806 nuclei and a mean of 2,361 genes per nucleus. From the five control samples we obtained 8,304 nuclei with a mean of 3,389 genes per nucleus (Fig 1a).

### Cell type composition

Using the snRNA-seq data, we performed an unbiased clustering using Seurat to identify and quantify the different cell types present in the injured brain tissue. We detected 15 different clusters and used canonical marker gene expression^17^ to identify the different cell types (Fig 1b-d). We identified three clusters of excitatory neurons and four clusters of inhibitory neurons, as well as clusters with oligodendrocytes, oligodendrocyte precursor cells (OPCs), astrocytes, endothelial cells, and microglia/macrophages (Fig 1c-d). All clusters were represented in both TBI and control samples, and the cellular composition was similar between samples obtained from temporal and frontal brain regions (Fig 1d and Suppl Fig 1b-c).

The most abundant cell type in the control samples was excitatory neurons, which accounted for almost 50% of the cells, followed by interneurons (20%) and astrocytes (13%) (Suppl Fig 1b). Oligodendrocytes made up 9% of the cells while microglia represented 2%. In the TBI samples we found a reduction in the proportion of excitatory neurons, which corresponded to 29% of the cells. In contrast, the proportions of both oligodendrocytes and microglia were higher (25% and 7%, respectively). The higher percentage of oligodendrocytes is likely explained by the contamination of white matter tissue in two of the TBI samples (Suppl Fig 1e) - which is difficult to control for in this clinical setting. However, the reduced numbers of excitatory neurons as well as the increase in microglia are likely to be due to the direct, acute consequences of the injury in line with the expected pathological outcome after severe TBI (Suppl. Fig 1b).

### OPCs and oligodendrocytes display an interferon response after acute TBI

The neuroinflammatory response is thought to be a key mechanism in the subsequent pathological processes following TBI. However, how this process starts and what cell types are involved in humans is unknown. To investigate transcriptional changes linked to inflammation, we analyzed cell-type-specific transcriptional alterations after TBI. We found that each cell type displayed a distinct transcriptional response to TBI. For example, in excitatory neurons we found that genes linked to synaptic functions, such as NPTX2 (padj 0.00; 2.45 log2FC. Wilcoxon Rank Sum test (FindMarkers, Seurat)) and HOMER1 (padj 2.78e-88; 2.1 log2FC), were upregulated: gene set enrichment analysis (GSEA) confirmed that genes linked to cell communication and synaptic signaling were dysregulated in excitatory neurons (Suppl Fig 2b). In microglia we found that genes linked to the cell cycle, such as MKI67 (padj 5.3e-6; 1.18 log2FC) and TOP2A (padj 1e-4; 1.02 log2FC), were upregulated, and GO analysis confirmed a transcriptional response linked to cell proliferation (Suppl Fig 2c). In line with this observation, a global analysis of cell-cycle-related genes (function ‘*CellCycleScoring*’, Seurat) confirmed that we detected microglial cells that were in a proliferative state in TBI samples and not in the controls (Suppl Fig 2f), which is in line with what is expected after TBI. No evidence of proliferation was detected in any other cell types after TBI. Thus, these results demonstrate that we detected neuronal dysfunction and the initiation of a microglia response upon TBI using the snRNA-seq approach.

**Figure 2.**
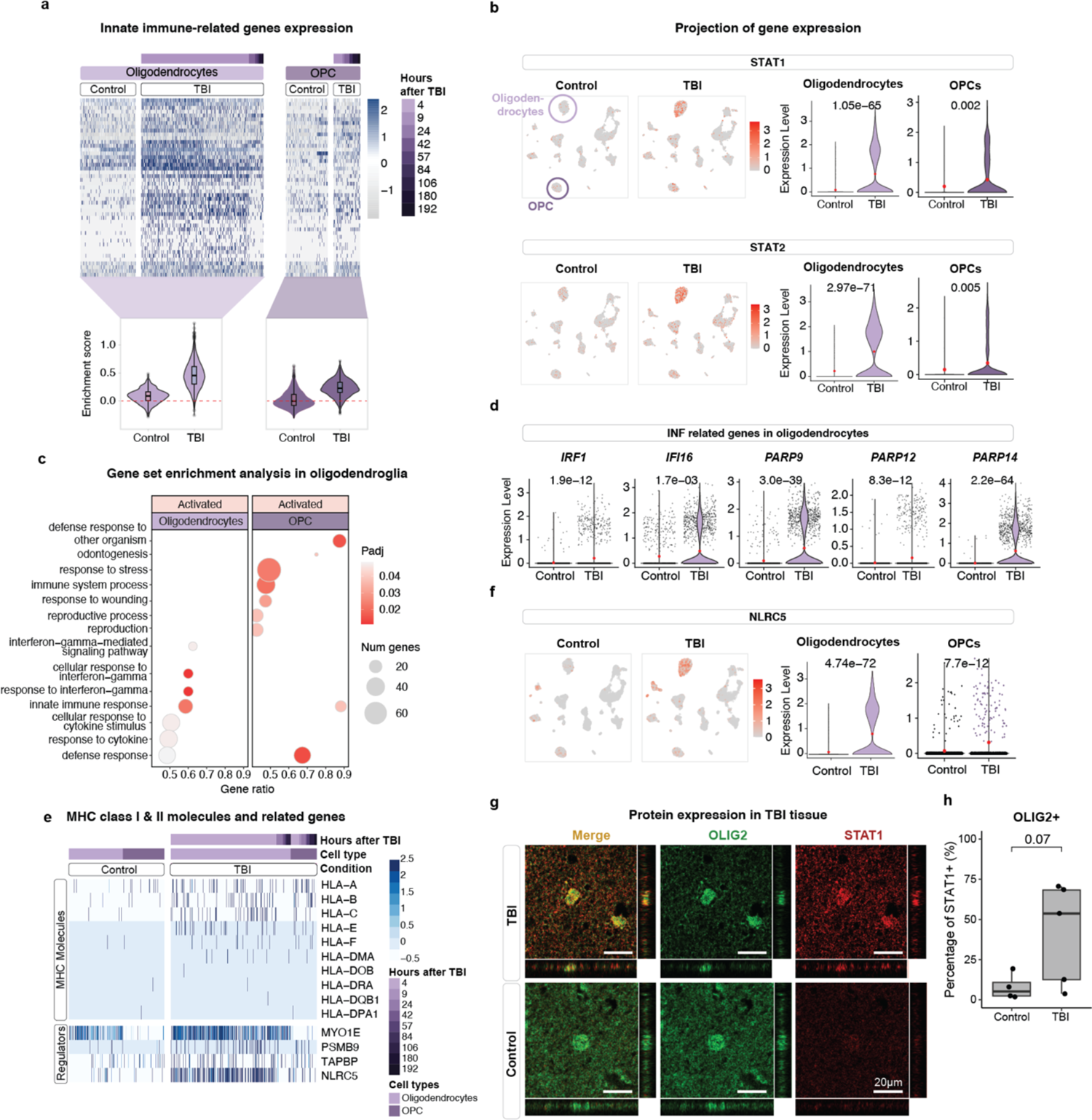
a) Top: Heatmap of oligodendroglia expression of differentially expressed genes (padj < 0.01; log2FC > 0.05) related to an innate immune response. Bottom: Enrichment scores per condition of the genes shown in heatmap (AddModuleScore, Seurat). B) Top panel showing STAT1 expression projected onto control and TBI UMAPs (left); violin plots showing expression per condition in oligodendroglia (right; Wilcoxon Rank Sum test (FindMarkers, Seurat)). Bottom panel showing STAT2 expression projected onto control and TBI UMAPs (left); violin plots showing expression per condition in oligodendroglia (right; Wilcoxon Rank Sum test (FindMarkers, Seurat)). c) Activated terms from GSEA of differentially expressed genes (TBI vs control; padj < 0.01, Wilcoxon Rank Sum test (FindMarkers, Seurat)) in oligodendroglia (biological-process ontology). c) Violin plots showing the expression of selected genes related to defense, interferon response, or inflammation in oligodendrocytes. e) Expression of detected molecules (top) of MHC class I and class II and related genes found to be significantly upregulated (bottom) in oligodendroglia. f) Expression of the key regulator of MHC class II molecules, NLRC5, projected onto control and TBI UMAPs (left); violin plot showing NLRC5 expression per condition in oligodendroglia (right; Wilcoxon Rank Sum test (FindMarkers, Seurat)). g) Immunohistochemical co-labelling of STAT1 (red) and OLIG2 (green). Confocal microscopy analysis revealed the presence of STAT1-expressing OLIG2-positve oligodendrocytes in TBI tissue. Scale bar represents 20μm. h) Automated microscopy quantification of the percentage of STAT1-positive cells among OLIG2-positive cells in TBI and control tissue (p = 0.073, Students t-test, n=5 TBI, n=4 ctrl).

When investigating transcriptomic changes in the other cell types, we found that OPCs and mature oligodendrocytes displayed a unique transcriptional response linked to neuroinflammation following severe TBI. In both cell types, we found that many genes involved in innate immunity and an interferon response, including STAT1 and STAT2, were activated (Fig 2a-b). GSEA confirmed that upon TBI, genes related to terms such as innate immune response and defense response were significantly enriched among the upregulated genes in both OPCs and mature oligodendrocytes (Fig 2c). Particularly in mature oligodendrocytes, we also found the activation of terms related to a response to interferon-gamma and cytokine stimulus (Fig 2c) showing clear upregulation of interferon regulatory factor IRF1 (padj 1,88e-12; 0,64 log2FC), interferon induced IFI16 (padj 0.001; 0,53 log2FC), and several PARP genes which respond to DNA damage and have different antiviral properties (PARP9 (padj 3,02e-39; 1,13 log2FC), PARP12 (padj 8,26e-12; 0.44 log2FC), and PARP14 (padj 2,22e-64; 1,63 log2FC)) (Fig 2d) ^23, 24^. Genes related to this response were found to be especially enriched in oligodendrocytes of samples collected a short time after the injury (up to 4 hours) (Suppl Fig 2a). These observations indicate that oligodendroglia play a key role in the initiation of inflammation following TBI via the activation of an interferon response.

Many of the genes upregulated in TBI oligodendroglia were MHC class I genes as well as regulators of class I and class II (Fig 2e-f and Suppl Fig 3b-c). For example, TBI oligodendroglia was found to have a significantly higher expression of NLRC5 (oligodendrocytes padj 4,74e-72; 1,55 log2FC. OPCs padj 7,65e-12; 0,6 log2FC) (Fig 2f), the major regulator of genes in the MHC class II^25^, as well as expressing other related genes such as PSMB9 (oligodendrocytes padj 9.65e-09, log2FC 0.49) and TAPBP (oligodendrocytes padj 5.87e-05, log2FC 0.41) which process MHC class I molecules and peptides, and MYO1E (oligodendrocytes padj 1.43e-12; log2FC 0.72) which regulates antigen presentation of MHC class II molecules ^26^ (Fig 2e). When we performed a similar analysis on microglia and astrocytes, we found no evidence of an innate or interferon response, nor the upregulation of MHC molecules (Suppl. Fig 3a-c).

**Figure 3.**
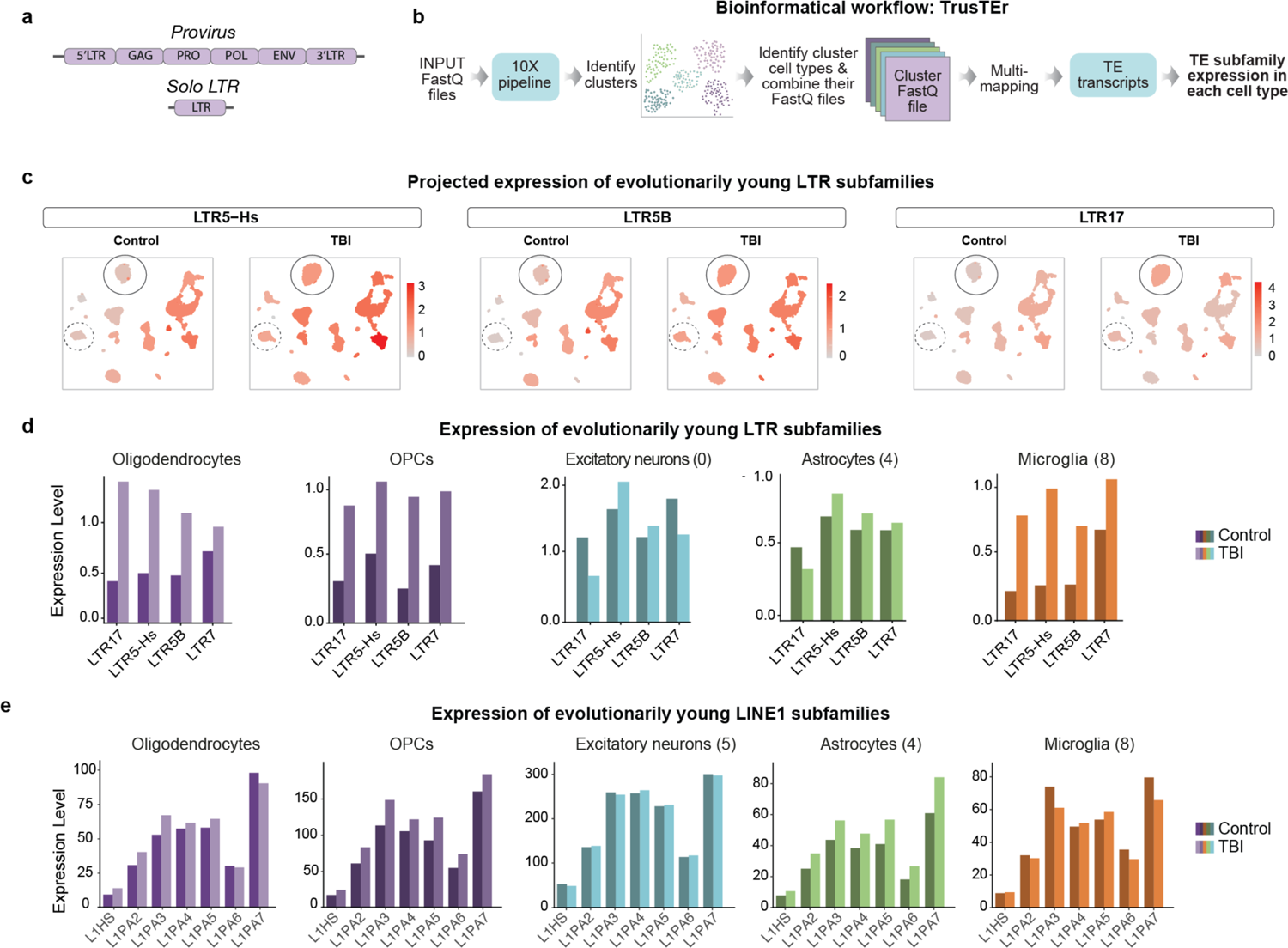
a) Schematic of the structure of an ERV provirus (top) and solo LTR (bottom). b) Schematic of bioinformatical approach to quantify TE subfamilies per cell cluster. c) Split UMAPs (control and TBI) with projected expression of LTR subfamilies per cluster. Oligodendrocytes circled, and OPCs circled in dotted line. d) Expression of evolutionary young LTR subfamilies in oligodendrocytes and OPCs. e) Expression of evolutionary young L1 subfamilies in oligodendrocytes and OPCs.

To confirm an activation of the interferon response at the protein level, we performed immunohistochemistry (IHC) in TBI and control tissue for STAT1 in combination with OLIG2, which specifically marks oligodendrocytes (Fig 2g). Confocal microscopy confirmed numerous STAT1+/OLIG2+ cells in TBI tissue. We used automated microscopy analysis to quantify the number of STAT1 expressing OLIG2-postive cells. While control tissue exhibited sparse STAT1 expression in oligodendrocytes, we observed an induction of STAT1 expression in oligodendrocytes in TBI tissue, although the signal was variable among individuals due to the technical challenges to perform robust IHC analysis on the severely damaged TBI tissue (Fig 2h, 41% in TBI vs. 8% in ctrl, p-value = 0.073 Students t-test). Taken together, these results indicate that oligodendroglia undergo a unique transcriptional response following TBI, which activates an interferon response, and turns-on MHC-related genes – hereby transforming to an immune-like cell state.

### Activation of endogenous retroviruses in oligodendrocytes after TBI

The transcriptional response in OPCs and mature oligodendrocytes after TBI is reminiscent to that which occurs after viral infection. In this respect, the transcriptional activation of ERVs (Fig 3a) has been linked to an interferon response. The aberrant expression of ERVs results in the formation of double-stranded RNAs, reverse-transcribed DNA molecules, and ERV-derived peptides that induce a “viral mimicry”, where cells respond as though infected and this triggers or boosts inflammation ^18–20^.

To investigate if ERVs, or any other TEs, were activated following TBI, we used an in-house bioinformatical pipeline ^17^ allowing the analysis of ERV and TE expression from our 10X single-nuclei RNA-seq data set (Fig 3b). In brief, this method uses the cell clusters determined based on the gene expression. Then, by back-tracing the reads from cells forming each cluster, it is possible to analyze the expression of ERVs and other TEs using the specialized software *TEtranscripts* ^27^ in distinct cell populations. This approach greatly increases the sensitivity of the analysis and enables quantitative estimation of ERV expression at single-cell-type resolution.

When using this bioinformatical approach, we found that several ERV (long terminal repeat – LTR) subfamilies were transcriptionally activated in oligodendrocytes and OPCs. This response was especially noticeable in ERV subfamilies such as HERV-K (LTR5-Hs & LTR5B, fold change of 2.34-2.7 in oligodendrocytes, 2-3.7 in OPCs), HERV-W (LTR17, fold change 3.41 in oligodendrocytes, 2.8 in OPCs) and HERV-H (LTR7 fold change 1.34 in oligodendrocytes, 2.3 in OPCs) which are evolutionary young ERVs found specifically in primates (Fig 3c-d). We found no transcriptional activation of other families of transposable elements such as LINE-1s (Fig 3e), and no evidence for the activation of ERVs or other TEs in other cell types, except for microglia which also displayed some evidence of ERV activation (Suppl Fig 4a-b).

**Figure 4.**
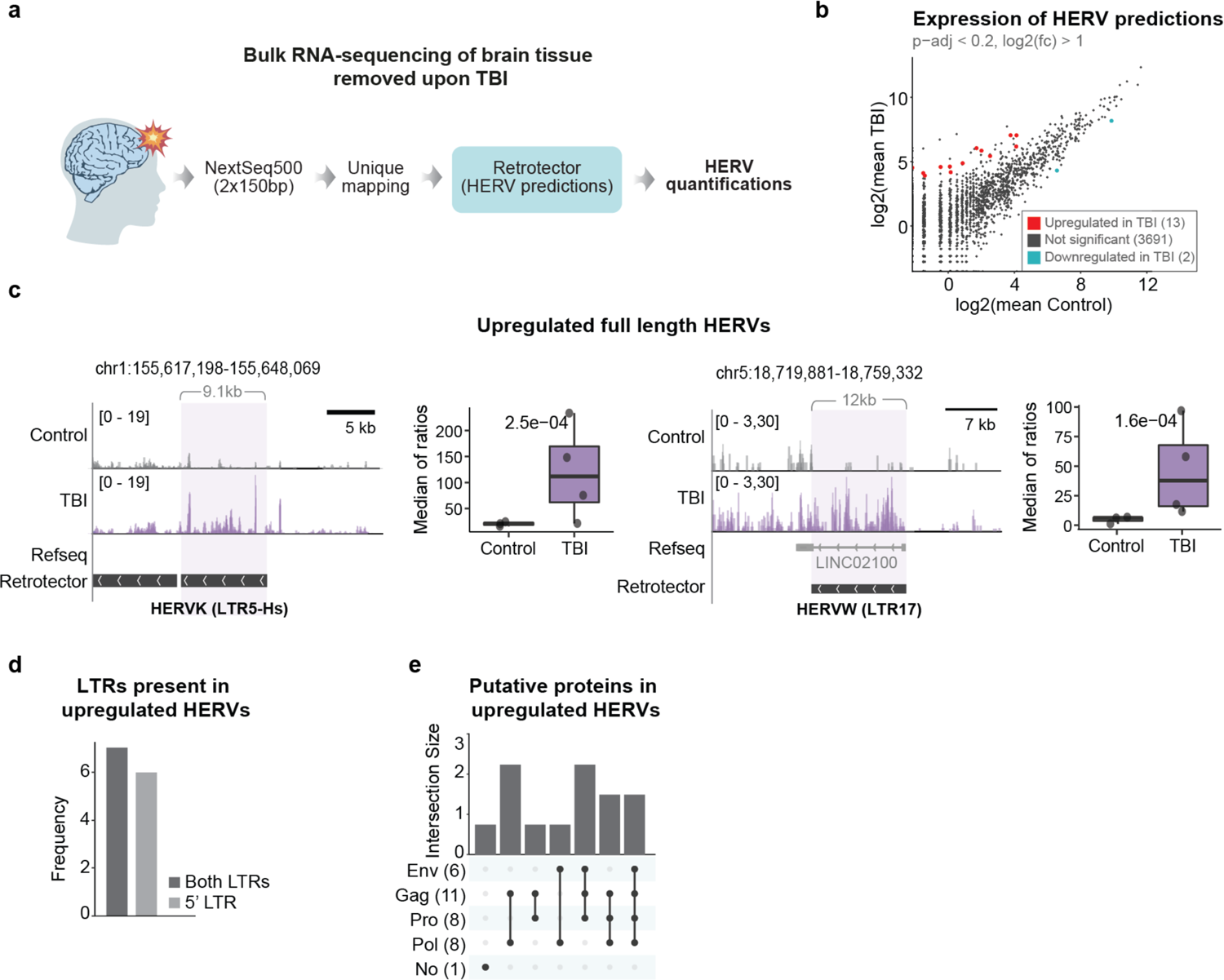
a) Schematic for bulk RNA-seq and bioinformatical approach. b) Scatter plot showing mean expression per condition and statistical analysis results for differential expression of all ERV predictions (DESeq2; p-value < 0.05, log2FC > 1 highlighted in red; p-value < 0.05 highlighted in blue). c) Tracks per condition with two examples of upregulated HERVs (left - HERVK, right - HERVW). Box plots showing quantification of the respective element (normalization by median of ratios and statistical test performed by DESeq2). d) Bar plots showing the number of upregulated ERV predictions with both LTRs or only the 5’ LTR. e) Upset plot showing putative proteins contained in the upregulated ERVs.

Many of the ERV families that were upregulated in TBI oligodendrocytes include provirus insertions with the potential to be transcribed and translated to produce ERV-derived peptides (top panel Fig 3a). Such ERVs have previously been linked to viral mimicry and the induction of an interferon response ^19^. However, since 10X libraries display a 3’ bias, it is not possible to distinguish between ERVs and solo LTR fragments that are present in large numbers in the human genome due to recombination events between the two endogenous provirus LTRs during primate evolution (Fig 3a). Thus, to investigate which unique loci the upregulated ERV expression in TBI oligodendrocytes originates from, we performed deep 2×150 bp paired-end, strand-specific bulk RNA-seq of four of the TBI samples with a high composition of oligodendroglia, as well as the control samples (Fig 4a and Suppl Fig 1a). Using a unique-mapping approach, we quantified ERV predictions identified using RetroTector ^28^ and found 13 significantly upregulated ERV loci in the TBI samples (*DESeq2*, p-value < 0.05, log2FC > 1) (Fig 4b, Suppl Fig 5a) which range in length from 6.2 kbp to 13.5 kbp, where transcription was found across the ERV sequence (Fig 4c). Notably, using our snRNAseq dataset, we were able to verify that some of the highly expressed, upregulated ERV loci were only expressed in oligodendroglia (Suppl Fig 4b). Six of these upregulated ERVs retained only the 5’ LTR, while the remaining seven retained both the 5’ and 3’ LTRs (Fig 4d). Furthermore, the predicted structures of these elements show that 12 out of the 13 upregulated ERVs still contain at least one ORF of the ERV-genes (*gag, pro, pol* and *env*) (Fig 4e). These data demonstrate that ERVs, with the potential to induce viral mimicry, are transcriptionally activated in oligodendrocytes within hours after TBI.

**Figure 5.**
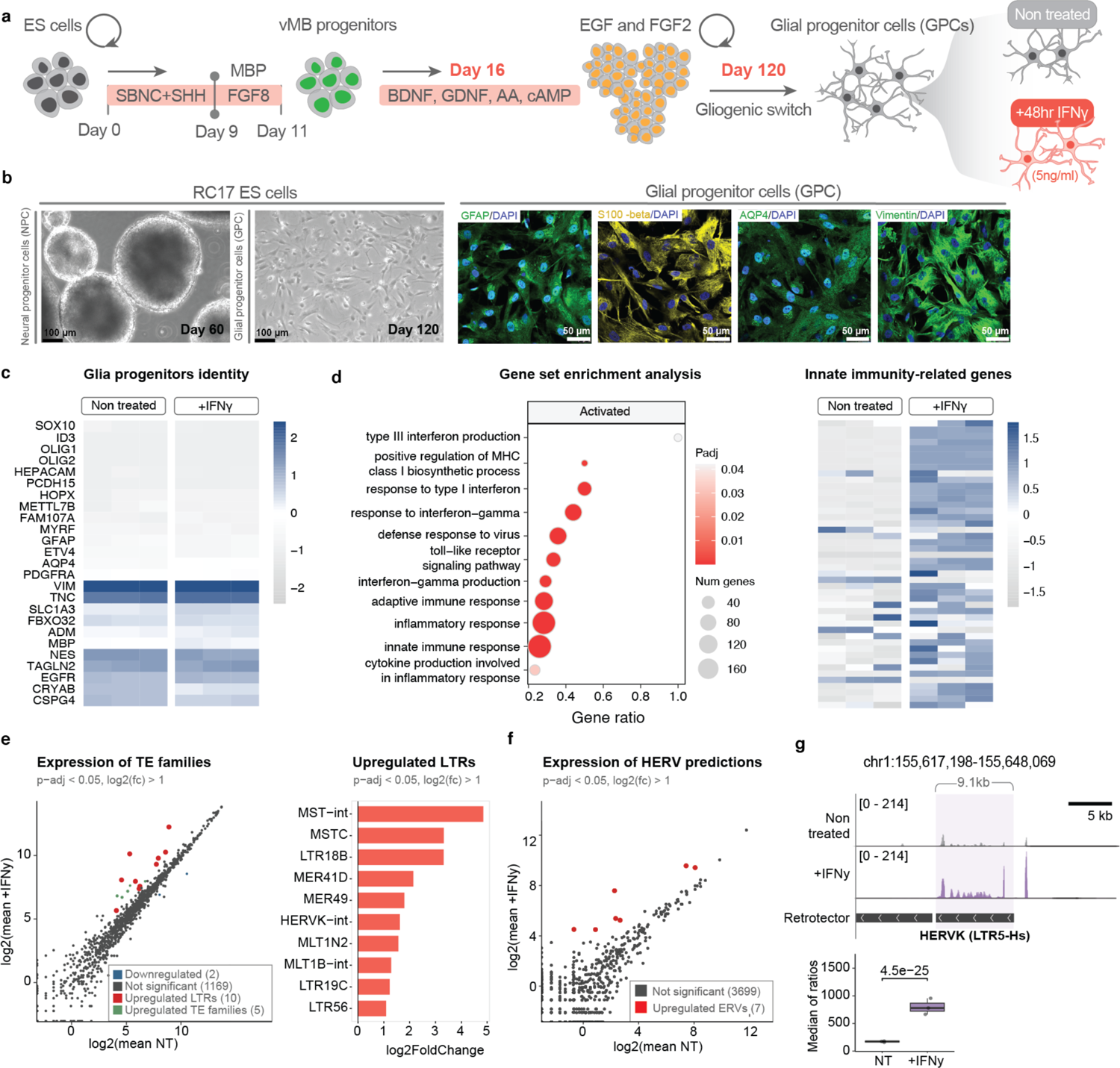
a) Schematic representation of differentiation protocol for deriving glial progenitor cells from hESCs and experimental approach for IFNγ treatment. b) Left: Bright-field images of hESC-derived vMB neural progenitor cells at day 60 maintained in a proliferation medium containing EGF and FGF2 (black scale bar: 100μm). Right: Immunocytochemistry of GPCs at day 120 of glia (GFAP, S100-beta, AQP4, Vimentin) and nuclear (DAPI) markers (white scale bar: 50μm). c) Canonical gene markers to validate glia progenitor cell identity. d) Left: Selection of activated GO terms upon IFNγ treatment. Right: Expression of innate immune related genes which were found to be upregulated in TBI oligodendroglia (as shown in *Figure 2a*). e) Left: Scatter plot showing mean expression per condition and statistical analysis results for differential expression of TE subfamilies (non-treated [NT] vs +IFNγ). Upregulated LTR subfamilies coloured in red, other upregulated TE subfamilies (non-LTRs) coloured in green (DESeq2; padj < 0.05; log2FC > 1). Downregulated subfamilies coloured in blue (DESeq2; padj < 0.05; log2FC < −1). Right: Bar plot showing log2FoldChange of upregulated LTR subfamilies (DESeq2; padj < 0.05; log2FC > 1). f) Scatter plot showing mean expression per condition and statistical analysis results for differential expression of all ERV predictions. DESeq2; p-value < 0.05, log2FC > 1 highlighted in red; p-value < 0.05 highlighted in blue. g) Tracks per condition showing an upregulated HERV-K (as shown in *Figure 4c*). Boxplot showing quantification of the element (normalization by median of ratios and statistical test performed by DESeq2).

### Interferon treatment results in transcriptional activation of ERVs in human glial progenitor cells

The induction of an interferon response has been associated with the transcriptional activation of ERVs, which is thought to play a part in boosting the interferon response^29^. To investigate a mechanistic link between interferon activation and ERV expression we decided to perform *in vitro* experiments in human glial progenitor cells (hGPCs).

We differentiated human embryonic stem cells (hESCs) into hGPCs using a 135-day differentiation protocol (Fig 5a, see methods section for details). The differentiated cells displayed a morphology of human glial cells and expressed marker genes related to this cell state, including many genes expressed in OPCs *in vivo* (Fig 5b-c). The hGPCs were treated with interferon gamma (IFNγ, 5ng/ml) for 48 hours before harvested for 2×150 bp paired-end, strand-specific bulk RNA-seq analysis.

When investigating changes in gene expression profile upon IFNγ stimulation of hGPCs, we found a distinct and robust transcriptional response. Genes linked to IFN signaling and innate immune activation were highly upregulated in the IFNγ treated hGPCs (Fig 5d). Thus, these observations confirm that hES-derived GPCs respond to interferon gamma treatment by transcriptional activation of its downstream targets (IFN-stimulated genes).

To quantify ERV expression upon IFN-y treatment in hGPCs we used two different bioinformatic methodologies. First, we allowed reads to map to different locations (multi-mapping) and used the *TEtranscripts* software ^30^ in multi-mode to best assign these reads and quantified the expression of different ERV subfamilies. Second, we discarded all ambiguously mapping reads and only quantified those that map uniquely and quantified the expression of unique ERV loci. The *TE-transcripts* approach revealed that several ERV subfamilies were robustly upregulated upon IFNγ treatment (Fig 5e). The upregulated ERV families included for example HERV-K, which are evolutionarily young ERVs with the capacity to trigger viral mimicry and that we also found to be upregulated in the TBI samples. Notably, we found no activation of other TEs, such as LINE-1s, indicating that the response to IFNγ treatment results in a transcriptional activation response specific to ERVs. The analysis of individual ERV loci identified several proviruses that were upregulated (Fig 5f). One of the most robustly upregulated proviral insertions was the same HERV-K provirus on chromosome 1 that we also found to be upregulated in the TBI samples (Fig 5g – compare with Fig 4c). Taken together, this experiment demonstrates that the activation of an interferon response in hGPCs results in transcriptional activation of ERVs, thus providing a mechanistic explanation to our observations of interferon activation and ERV-expression in TBI-tissue.

## Discussion

Neuroinflammation is a hallmark of acute and chronic neurodegenerative states, including TBI, and is therefore a promising route for urgently needed disease modifying therapies. However, information on how the neuroinflammatory response starts and then transforms to a chronic state is scarce, and the molecular events needed to initiate and maintain this process, and in which cell types, has not been established. This lack of basic understanding hampers the development of new treatment strategies. In this study, we used snRNA-seq to perform a detailed analysis of human tissue samples obtained in conjunction with acute surgery for TBI. Our results provide two unique insights into the start of a human neuroinflammatory response. First, we found that OPCs and oligodendrocytes play an unanticipated key role in this process by initiating an interferon response and adopting an immune-like cell state. Secondly, we found that the activation of an interferon response in OPCs and oligodendrocytes is mechanistically linked to the transcriptional activation of ERVs.

Neuronal cell death is common in acute, severe TBI due to hemorrhages, increased intracranial pressure, and energy metabolic failure. In addition, there is a distinct inflammatory response, associated with white matter abnormalities, that persists from the acute post-injury phase for many years post-injury ^2, 6, 31, 32^. Prior studies have established that there are numerous contributors to the neuroinflammatory response after TBI, including microglial activation, infiltration of systemic immune cells, and the subsequent release of pro-inflammatory cytokines ^33, 34^. Our observations indicate that oligodendroglia also seem to play an important role in the initiation of this process. Oligodendroglia are vulnerable to the TBI-induced neuroinflammation as well as to excitotoxicity and reactive oxygen species formation ^35, 36^, and loss of mature oligodendrocytes is observed at an early stage following rodent ^37–39^ and human TBI ^40^. Treatment with an anti-inflammatory antibody neutralizing interleukin-1β, a key pro-inflammatory cytokine, attenuates this post-injury loss of oligodendrocytes in rodent models ^9^. The trigger of an interferon response upon TBI has been reported before ^41, 42^, although the specific cell types involved in this process remained unknown. In the present study, we found that OPCs and oligodendrocytes undergo the activation of interferon response genes as well as genes related to MHC class I and class II. Some of these genes affect the expression, processing, and guidance of MHC molecules, which are crucial to immune cell functions. Our results suggest that, following TBI, oligodendroglia adopt a different cell state characterized by the expression of immune genes. These results are reminiscent of what has been observed in multiple sclerosis, where OPCs and oligodendrocytes initiate a similar transcriptional program and adopt different functional properties, such as cell phagocytosis ^14, 15^. Future investigations into the role of oligodendroglia in the inflammatory response, and the underlying mechanisms linked to their cell fate change, could lead to new treatment opportunities.

Almost 10% of the human genome is made up of ERVs, a consequence of their colonization of our germline throughout evolution ^43^. In adult tissues, including the brain, ERVs are normally transcriptionally silenced via epigenetic mechanisms, including DNA and histone methylation ^16^. However, there is emerging evidence, both from animal models and the analysis of human material, indicating that ERVs can be transcriptionally activated in the diseased brain and that this correlates with neuroinflammation. ERV expression has been found to be elevated in the cerebrospinal fluid and in *post-mortem* brain biopsies from patients with multiple sclerosis, amyotrophic lateral sclerosis, and Alzheimer’s and Parkinson’s disease ^16, 44–52^. In experimental drosophila or mouse models there is causative evidence linking upregulation of ERVs and other transposable elements to neuroinflammation and neurodegeneration ^17, 53^. However, accurate estimation of ERV expression is challenging: their repetitive nature complicates the bioinformatical analysis and many of these observations remain controversial. Furthermore, most of the clinical studies use end-stage *post-mortem* material where the inflammatory process has been ongoing for decades or are limited to the study of biofluids such as cerebrospinal fluid or plasma. These limitations have made it challenging to conclusively link ERV expression to the initiation of the neuroinflammatory response. Our results now provide direct evidence that ERV proviruses are transcriptionally activated at the start of human neuroinflammation. Notably, we find that ERVs are specifically activated in oligodendrocytes and OPCs, the cell types involved in the interferon response. Thus, our results provide direct clinical evidence linking the induction of an immune response and the transcriptional activation of ERVs. Our mechanistic *in vitro* modeling in human glia progenitor cultures demonstrates that the induction of an innate immune response causes a transcriptional activation of ERVs. Exactly what triggers the interferon response remains unclear. It may be due to the activation of ERVs via epigenetic mechanisms, but we cannot rule out other possible mechanisms such as the release of mitochondrial DNA into the cytosol or genomic DNA damage.

From our results it is also not possible to pinpoint the role ERVs have in driving and boosting the immune response following their transcriptional activation. However, there is growing awareness that the transcriptional activation of ERVs results in the activation of immune pathways ^18–20^. For example, the same HERV-K provirus on chromosome 1 (also known as *ERVK-7* ^54^) that we found upregulated in TBI tissue and IFNγ-treated hGPC (Fig 4c and 5g) was recently reported to trigger the production of ERV-reactive antibodies in human lung cancer patients through the expression if its envelope glycoprotein ^54^. In addition, ERV-derived peptides may contribute to protein aggregation, which is a process directly linked to chronic effects following TBI and which also may further enhance an inflammatory response ^55^.

In summary, we here describe a novel role for OPCs and oligodendrocytes in the initiation of an interferon response following acute severe human TBI. We describe the activation of an interferon response in oligodendroglia that is linked to the activation of regulatory genes of molecules present in immune cells. This transcriptional switch in OPCs and oligodendrocytes is mechanistically linked to the activation of ERVs. Our results could lead to entirely new treatment opportunities for acute TBI, as well as for chronic neurodegenerative disorders.

## Methods

### Ethical statement

All clinical and experimental research described herein was approved by the regional ethical review board (decision numbers 2005/103, 2008/303, 2009/89, and 2010/379). Written informed consent was obtained from the TBI patients’ closest relatives and from the patients themselves if they had sufficiently recovered from their injury at >6 months post injury.

### TBI and control cohorts

Detailed demographic and clinical characteristics are shown in Table 1. Twelve patients with severe TBI, defined as post-resuscitation Glasgow Coma Scale score ≤ 8, were included. The patients were >18 years old, and no patient had any other known neurological disorder or Down’s syndrome. All patients had been mechanically ventilated and sedated and continuous measurements of intracranial pressure (ICP) and cerebral perfusion pressure (CPP) were performed at the neurocritical care unit at Uppsala University Hospital ^56^.

The age (mean ± SD) of the TBI patients (10 males, 2 females) was 49.5 ± 18.2 years. The tissue samples were obtained from patients suffering severe, life-threatening focal TBI. Due to space-occupying brain swelling causing midline shift and compression of the basal cisterns, or due to markedly increased intracranial pressure (ICP) refractory to medical neurointensive care measures, contused brain regions (typically the injured part of a temporal or frontal lobe) were surgically removed between 4 hours and 8 days after injury ^21^. There were no complications associated with the surgical method. The patients needed prolonged neurocritical, neurosurgical, and then neurorehabilitative care. At the time of follow-up, one patient was deceased. All available patients’ outcomes are shown in Table 1.

### Sampling and preparation of brain tissue

Surgically removed brain tissues were immediately placed in a sterile pre-labeled container and subsequently stored at −80°C until analyzed. Half of the tissue was put in a routinely-used fixative, 4% buffered formalin (Histo-Lab Products AB, Gothenburg, Sweden, catalogue no. 02176). The samples were fixed for 24-72 h and then paraffin-embedded and processed by hardware Tissue tek VIP (Sakura, CA, USA). Small samples were taken from the fresh-frozen contused brain tissue for snRNA-seq.

### Immunohistochemistry

Immunohistochemistry was performed on microtome sections according to previously published protocols^17^.

For OLIG2 and STAT1 two sets of antibodies were used with similar results. OLIG2 (R&D Systems, AF2418, 1:500), OLIG2 (Sigma-Aldrich, AB9610, 1:500), STAT1 (Abcam, ab109320, 1:500), STAT1 (Abcam, ab281999, 1:2000)

Confocal images were captures using a Leica TCS SP8 confocal laser-scanning microscope. The Operetta CLS (Perkin Elmer) instrument and Harmony analysis software (version 4.9.) were used for high-content screening and analysis. We identified the number of DAPI+ (Sigma-Aldrich, 1:1,000), STAT1+ and OLIG2+ cells and defined the number of cells with STAT1 staining. Tissue samples from five individuals with TBI and four individuals from non-neurological deaths were stained for DAPI, OLIG2 and STAT1. Images were acquired from 37-382 fields using the 20x objective. Valid nuclei were defined by DAPI staining based on intensity and area. We excluded DAPI cells which were clumped together or where the separation of nuclei by the software was not efficient enough by setting a maximum area and shape. STAT1+ and OLIG2+ cells were identified using a threshold considering the background of the staining.

### Statistical analysis

The percentage of STAT1 positive cells among OLIG2 positive cells was calculated only considering cells with valid DAPI staining as:

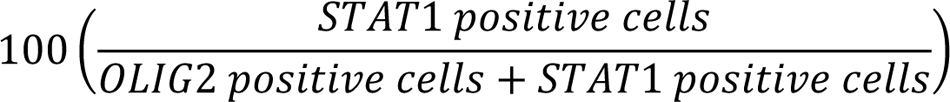

The data was normally distributed (p = 0.1397, Shapiro test for normality). We used Students t-test to identify the differences between the two groups. In the bar plot (Figure 2h), error lines indicate the standard error of the mean.

#### Single-nuclei RNA sequencing analysis

The FACS-based isolation of nuclei from frozen brain tissue was performed as previously described and loaded onto 10x Genomics Single Cell 3’ Chip ^17^. Sequencing library samples were multiplexed and sequenced on a Novaseq machine using a 150-cycle kit with the recommended read length from 10x Genomics. Raw base calls were demultiplexed to obtained sample-specific FastQ files and reads were aligned to GRCh38 genome assembly using the Cell Ranger pipeline (10x Genomics, Cellranger count v5; RRID:SCR_017344) with default parameters (--include-introns was used for nuclei mapping). The resulting matrix files were used for further downstream analysis. Seurat (version 3.1.1; RRID:SCR_007322) and R (version 3.4) were used for bioinformatics analysis.

Cells which have lower than three mean absolute deviation from the median number of reads present in the sample were removed (R function *scater::isOutlier,* parameters *type=*”both”*, nmads=3, log=TRUE)*. Given the difference between quality of TBI and control tissue, a further cut off at least 1,000 detected genes per cell was implemented to ensure that only high quality cells were retained for downstream analysis. Data was log-normalized to reduce sequencing depth variability (LogNormalize, Seurat). All samples were merged to generate integrated UMAP using Harmony integration (RunHarmony, ‘sample’, Seurat)^57^. For visualization and clustering, manifolds were calculated using UMAP methods (*RunUMAP,* Seurat) and 20 precomputed principal components and the shared nearest neighbor algorithm modularity optimization-based clustering algorithm (FindClusters, Seurat) with a resolution of 0.1. The clusters obtained were annotated using canonical marker gene expression. Differential gene expression between TBI and control samples across respective cell types was carried out using the Seurat function FindMarkers (Wilcoxon Rank Sum test, padj < 0.01). Cell-cycle scores were computed using the Seurat function CellCycleScoring. Gene ontology overrepresentation analysis was performed using the enrichGO function in the clusterProfiler R package using differentially-expressed genes in each cell type (padj < 0.01) ^58^. Enrichment scores for genes related to selected immune-related GO terms were calculated using the AddModuleScore function from Seurat using five features as controls (ctrl = 5).

#### TE quantification per cell cluster

To ease the usage of the workflow we made use of, we created a Python package named trusTEr (version 0.1.1; doi:10.5281/zenodo.7589548) to process single-cell RNA-seq data to quantify TE expression per cluster of cells. The package consists of three classes (Experiment, Sample, and Cluster) for the abstraction of the projects, and a module (jobHandler.py) to handle SLURM jobs. For more detailed information about how the classes are related, please refer to https://raquelgarza.github.io/truster/. The workflow has the following steps:

1. Extraction of reads from samples’ BAM files (tsv_to_bam() in class Cluster). After the clustering formation, trusTEr expects one text file per cluster containing all barcodes of a sample in a cluster (done by default by get_clusters() and merge_clusters()). It will extract the given barcodes from the sample’s BAM file (outs/possorted_genome_bam.bam output from Cell Ranger count) using the subset-bam software from 10x Genomics (version 1.0; RRID:SCR_023216). Outputs a BAM file per cluster containing all alignments from the cells in the cluster.
2. Filter duplicates (filter_UMIs() in class Cluster). Given the number of PCR duplicates we could be carrying in these BAM files, we wrote a Python script (filterUMIs) to filter most of these out. The filter_UMIs function keeps reads with unique combinations of cell barcodes, UMI, and sequence, so that only unique molecules are kept.
3. Convert to FastQ (bam_to_fastq() in class Cluster). Using bamtofastq from 10x Genomics (version 1.2.0; RRID: SCR_023215), the previous BAM files are output as fastQ files.
4. Concatenate lanes (concatenate_lanes() in class Cluster). Concatenates the different lanes from the same library as output from bamtofastq. This step outputs one fastQ file per cluster.
5. The quantification was performed in groups of samples (TBI and control groups) to increase statistical power. This step concatenates the fastQ files of the samples’ clusters within a single group (see the groups parameters in the process_clusters function in the Experiment class, or in each of the steps’ corresponding functions).
6. Map cluster (map_cluster() in class Cluster). Mapping the reads to a reference genome using STAR aligner (version 2.7.9a; RRID:SCR_004463) ^59^. For subfamily quantification (default in the map_cluster function) –outFilterMultimapNmax 100, -- winAnchorMultimapNmax 200 were used. To quantify individual elements (argument unique set to True), unique mapping was performed using –outFilterMultimapNmax 1 and –outFilterMismatchNoverLmax 0.03. Visualization of tracks per cell type (Suppl Fig 5b) was performed using the uniquely mapped reads per cluster, grouped by condition. These were filtered by strand using deeptools bamCoverage (version 2.4.3; RRID:SCR_016366)^60^ with --filterRNAstrand set to “forward” to get reverse transcription, and “reverse” to get forward transcription – as bamCoverage assumes a dUTP-based library preparation. Signal was normalized using a scale factor (--scaleFactor) of 1e+7 divided by the number of cells in the cluster. Tracks of cell types with more than one cluster were overlayed in the Integrative Genomics Viewer (IGV) (version 2.11.1; RRID:SCR_011793)^61^.
7. TE count (TE_count() in class Cluster). TE quantification of the BAM files produced for each cluster. For the purposes of this paper, we used TEcount from the TEToolkit (version 2.0.3; RRID:SCR_023208) with the curated TE GTF for hg38 provided by the authors (-- TE), gencode version 36 as the gene GTF (--GTF). We ran it in multi mode (--mode multi) as forward stranded (--stranded yes) ^30^.
8. Normalization of TE counts. TE quantification is normalized by cluster size (number of cells in a cluster), and it’s stored within the Seurat object. The matrices output and the Seurat assay only includes the normalized TE subfamily information output by TEcount.

#### Bulk RNA sequencing analysis

Total RNA was isolated from nuclei using the Rneasy Mini Kit (Qiagen). Libraries were generated using Illumina TruSeq Stranded mRNA library prep kit (poly-A selection) and were sequenced on an Illumina NextSeq500 machine (paired-end 2×150 bp).

Reads were mapped using STAR (2.6.0c; RRID:SCR_004463) ^59^ with GRCh38.p13 as the genome index, and gencode version 38 to guide the mapping (--sjdbGTFfile). To avoid ambiguous reads due to the quantification of ERVs, the number of loci allowed for a read to map (-- outFilterMultimapNmax) was set to 1, and the number of mismatches allowed (-- outFilterMismatchNoverLmax) was set to 3%.

The RetroTector software ^28^, (https://github.com/PatricJernLab/RetroTector) was used to mine the human genome (version GRCh38/hg38 downloaded from https://hgdownload.soe.ucsc.edu/) for ERVs as previously described.^62^ Predicted positions and structures for ERVs scoring 300 and above are summarized in Suppl. Tables 1 and 2. Using the GTF version of the output file, read were quantified using featureCounts (Subread version 1.6.3; RRID:SCR_012919) ^63^, forcing matching strandness of the reads to the features being quantified (-s 2).

Read count matrices were then input to DESeq2 (version 1.28.1; RRID:SCR_015687) and fold changes shrunk using DESeq2::lfcShrink ^64^. Volcano plots were visualized using the R package EnhancedVolcano (version 1.14.0) (https://github.com/kevinblighe/EnhancedVolcano). For further details, please refer to the corresponding Rmarkdown (TBI_bulk.Rmd) on github.

#### Generation of glial progenitors from hES cells

Human Embryonic stem cells (RC17) were maintained in IPS-brew on laminin 521 (0.5 μg/cm2) coated plates. Cells were passaged every 5 days with 0.5 mM EDTA, followed by seeding at a density of 2,500 cells per cm2 with ROCK inhibitor (10 μM Y-27632-only first 24 h after plating). RC17 ES cells were differentiated towards ventral midbrain (vMB) fate based on a previously published protocol (Nolbrant et al, 2017). Post quality control for ventral midbrain specification, day 16 progenitors were maintained in a Neurobasal medium containing NB-21 supplement without vitamin A (1:500), penicillin/streptomycin (1:1000), l-glutamine (1:1000), NEAA (1:1000), FGF2 (20 ng ml−1) and EGF (100 ng ml−1) in suspension on a non-adherent flask. vMB progenitors self-organise and form homogenous embryoid bodies (EBs) in suspension. EBs were dissociated every 18-20 days and maintained on a non-adherent flask for another 100 days for cells to undergo a gliogenic switch. Post 100 days of differentiation, cells were dissociated into single cells with accutase and plated on laminin 521 (2 μg/cm2) coated plates for terminal differentiation medium (Neurobasal medium containing NB-21 supplement without vitamin A (1:500), penicillin/streptomycin (1:1000), l-glutamine (1:1000), NEAA (1:1000) and CNTF (50 ng ml−1)) for glial progenitors.

#### Immunocytochemistry

Cells were fixed in 4% PFA for 20 min, and post-fixation cells were washed with 1XPBS and incubated in blocking buffer/ permeabilization buffer (Triton X-100) for 1 hour. Cells were incubated with primary antibodies overnight in PBS. Antibodies: GFAP (DAKO, Z0334, 1:500), AQP4 (Sigma, HPA014784, 1:500), s100-Beta (Sigma, S2532, 1:500) and Vimentin (Millipore, AB5733, 1:500). After incubation with secondary antibodies and DAPI, images were acquired in Lieca SP8 confocal microscope.

#### IFN-γ stimulation of glial progenitors

RC17-derived glial progenitor cells were plated on laminin 521 (2μg/cm2) coated plates in a terminal differentiation medium. Cells were matured for 14 days and stimulated with IFN-γ (5ng/ml) for 48hrs in the culture medium. Post stimulation, cells were washed twice with 1XPBS and detached with accutase and pellets were frozen down on dry ice for bulk-RNA seq.

### Data and code availability

Raw data of single-nuclei and bulk RNA-seq of the control post-mortem samples, and processed data of all samples can be found at the GEO repository GSE209552. Visualization, statistical analyses, and preprocessing pipelines, including QC filtering for the single nuclei RNAseq can be found at https://github.com/Jakobsson-Lab/TBI_Garza_2023. Code for the quantification of TEs per cluster from single nuclei RNAseq can can be found at https://github.com/raquelgarza/truster, documentation at https://raquelgarza.github.io/truster/.

## Acknowledgements

We would like to thank Roger A. Barker for critical input on the study and Alafuzoff, J. Johansson, U. Jarl, M Vejgården, B. Mattsson and A. Hammarberg for technical assistance. We are grateful to all members of the Jakobsson and Marklund labs. The work was supported by grants from the Swedish Research Council (2018-02694 to J.J., 2018-02500 to N.M., 2021-01740 to P.J.), the Swedish Brain Foundation (FO2019-0098 to J.J., FO2020-0147to NM), the Novo Nordisk Foundation (NNF21CC0073729 to AK) and the Swedish Government Initiative for Strategic Research Areas (MultiPark & StemTherapy).

## Author contributions

All of the authors took part in designing the study and interpreting the data. R.G., J.J., and N.M. conceived and designed the study. D.A., A.T., S.A.H, V.H., A.A., and M.J. performed the clinical and experimental research. R.G. and Y.S. performed the bioinformatic analyses. M.I., M.G.H., P.J., E.E and A.K. contributed reagents and expertise. R.G., J.J., and N.M. wrote the manuscript, and all authors reviewed the final version.

---

**Suppl. Figure 1.**
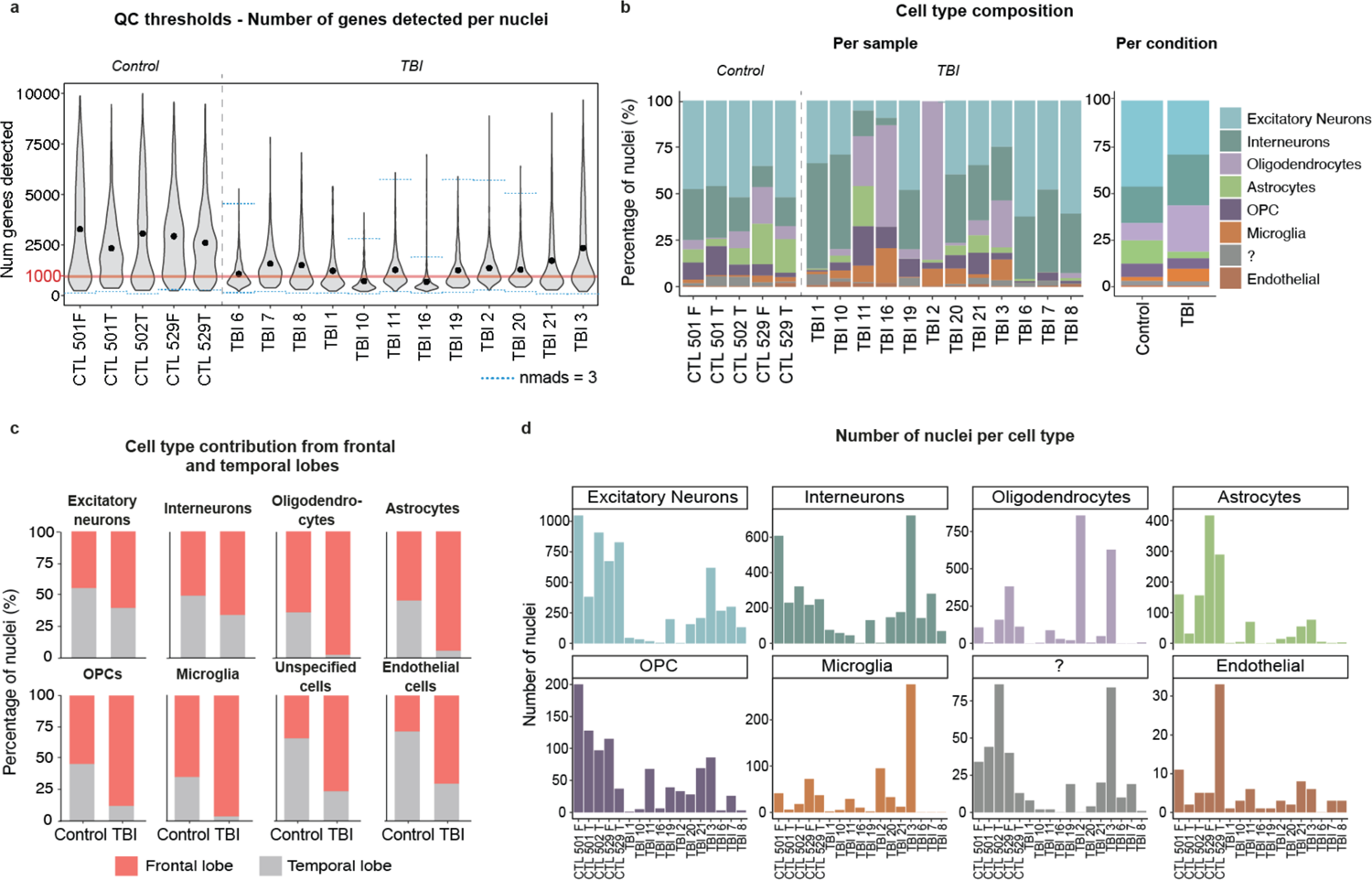
a) Quality control thresholds for number of genes detected. Blue dotted lines indicating thresholds of nuclei excluded as outliers (isOutlier, scater, nmads = 3; see methods). Nuclei below red line (1,000 genes detected) were excluded from downstream analysis. b) Cell-type ratio found in each sample and condition. c) Percentage of cells belonging to samples collected from the frontal or temporal lobe in each cell type. d) Number of nuclei per cell type in each sample.

**Suppl. Figure 2.**
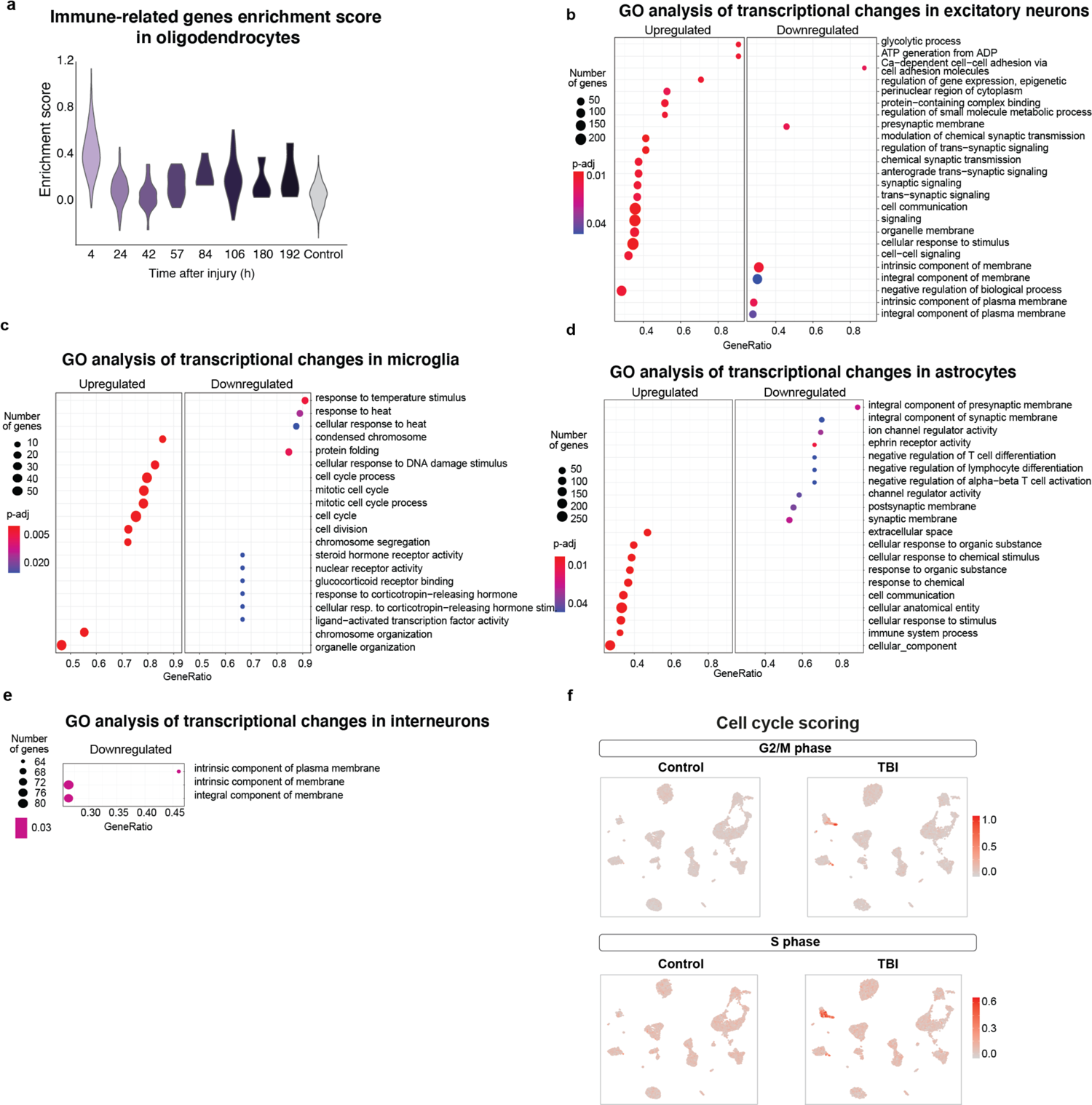
a) Violin plots showing enrichment scores of immune-related genes per time point of injury (GO terms: Interferon-gamma-mediated signaling pathway, Response to interferon-gamma, Cellular response to interferon-gamma, Innate immune response, Cellular-response to cytokine stimulus, Response to cytokine, and Defense response). b) Gene set enrichment analysis (GSEA) of differentially-expressed genes (padj < 0.01) found in excitatory neurons, c) microglia, d) astrocytes, and e) interneurons. f) Enrichment score of cell-cycle-related genes (top G2/M phase, bottom S phase).

**Suppl. Figure 3.**
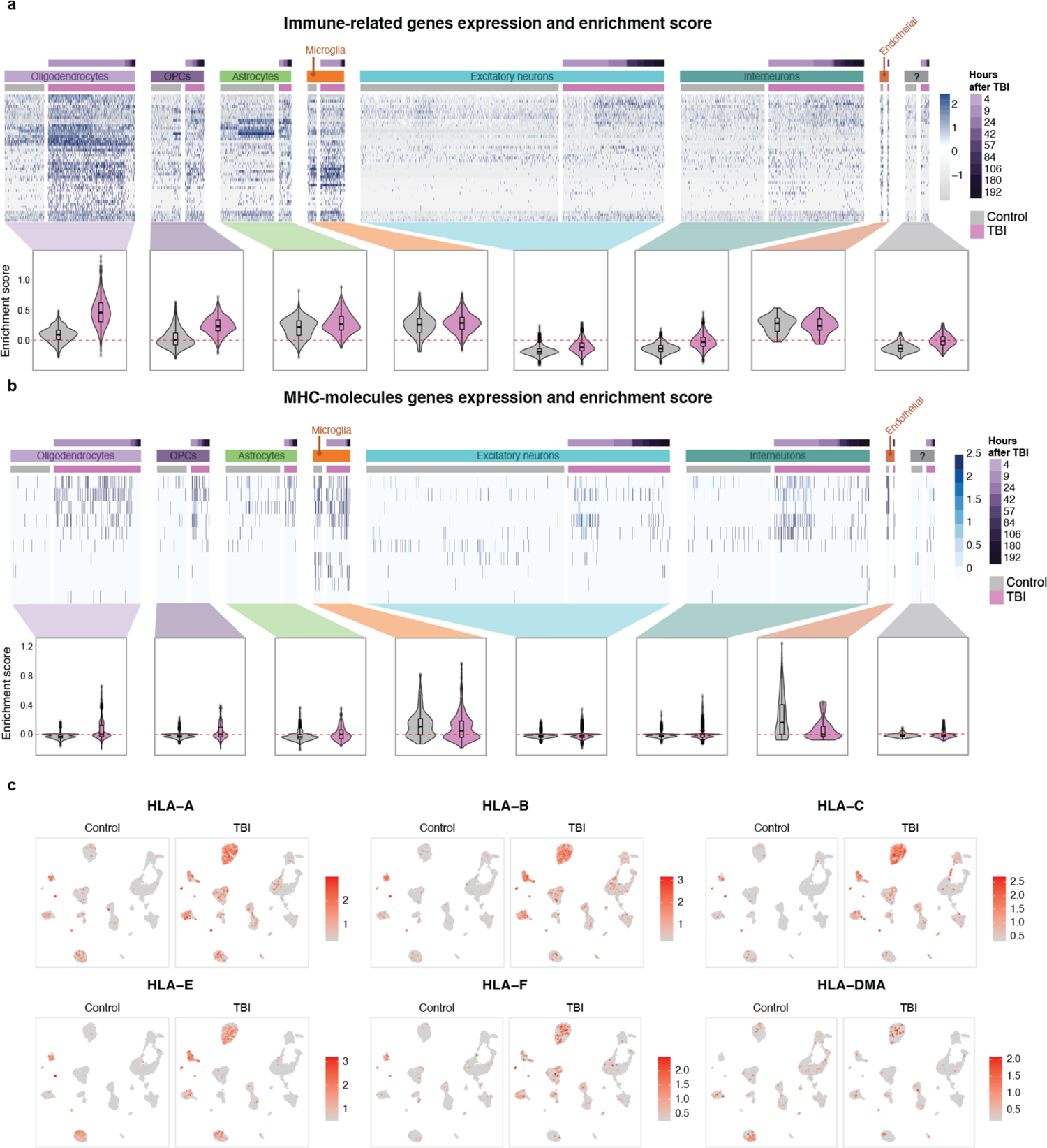
a) Heatmap showing expression of selected genes (upregulated genes in oligodendrocytes related to Interferon-gamma-mediated signaling pathway, Response to interferon-gamma, Cellular response to interferon-gamma, Innate immune response, Cellular-response to cytokine stimulus, Response to cytokine, and Defense response), as shown in *Figure 2a* and violin plots showing enrichment scores per cell type and condition (AddModuleScore, Seurat). b) Heatmap showing expression of detected genes encoding for MHC molecules, and violin plots showing enrichment scores per cell type and condition (AddModuleScore, Seurat). c) Expression of selected MHC molecules.

**Suppl. Figure 4.**
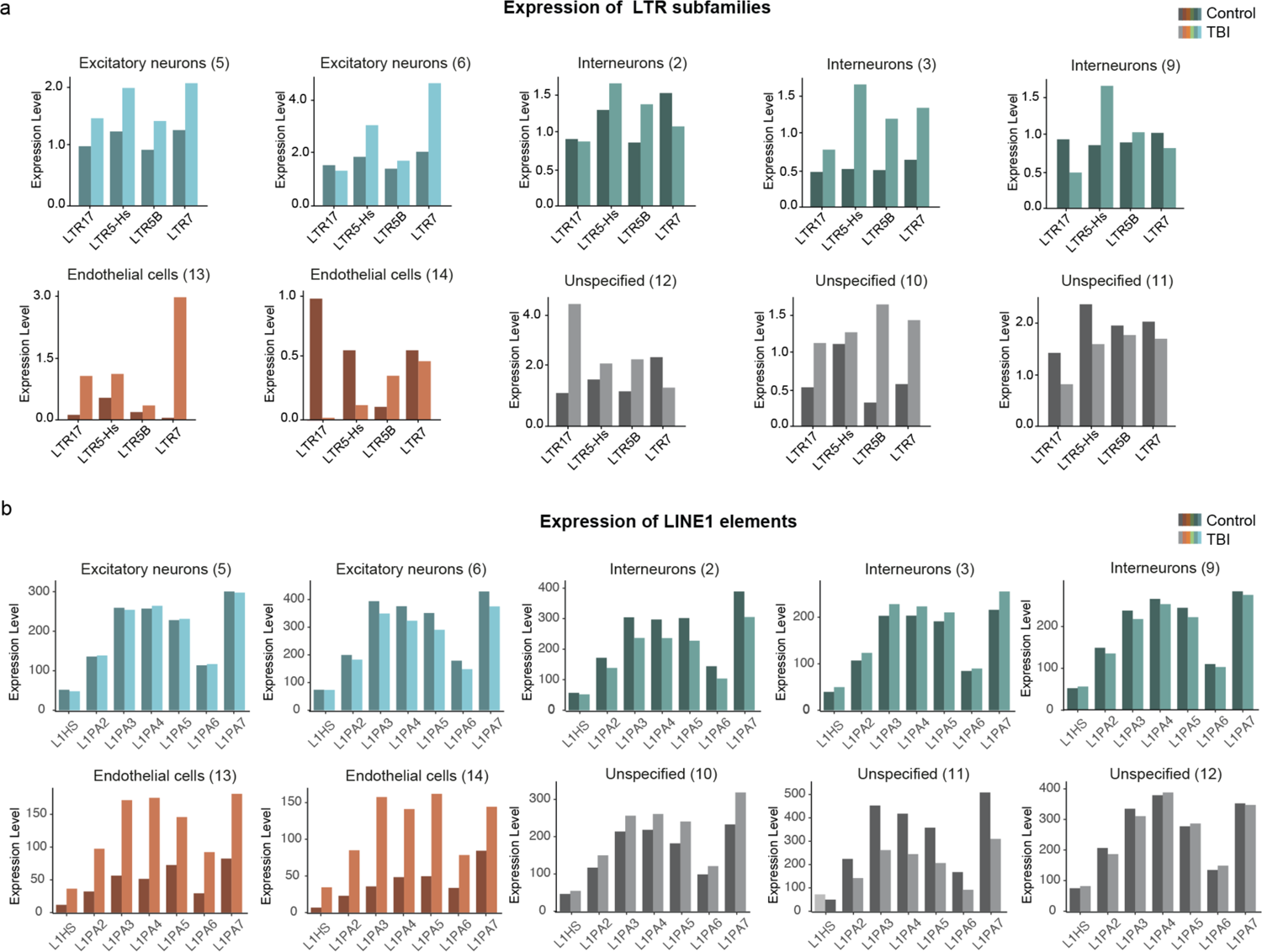
a) Expression of LTR subfamilies in the cell types not shown in *Figure 3d*. b) Expression of LINE1 subfamilies in the cell types not shown in *Figure 3e*.

**Suppl. Figure 5.**
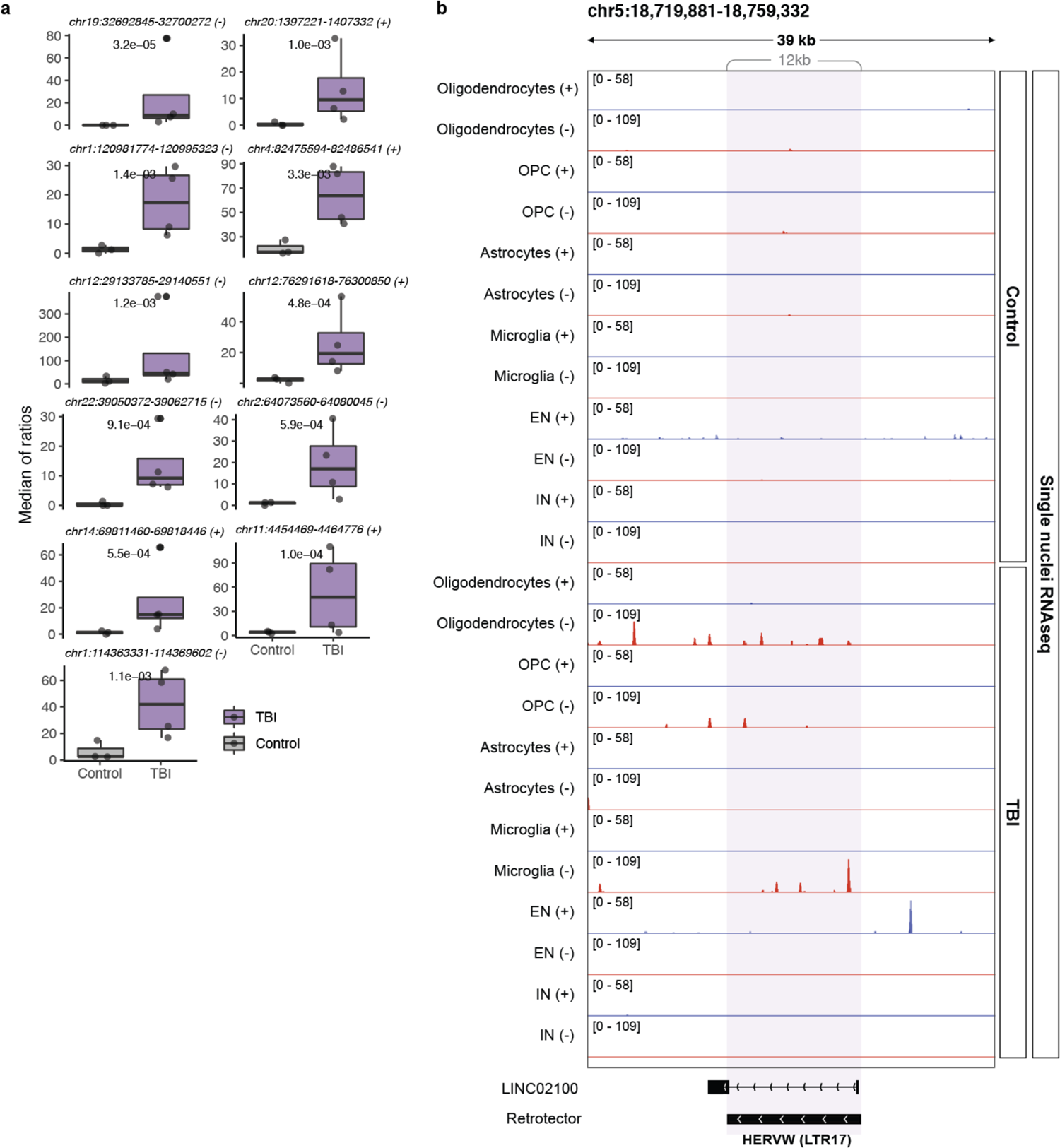
a) Boxplots showing normalized expression of upregulated ERVs per sample, grouped by condition (DESeq2 pvalue, unadjusted). b) Genome browser tracks showing an example of an upregulated HERV-W (*Figure 4c*). Cells were grouped by cluster and condition (TBI/Control) and uniquely mapped as pseudobulk. Tracks of clusters of same cell type are overlayed. Tracks marked as + (blue) or – (red) referring to forward and reverse transcription, respectively.

